# Competition among kin generates balancing selection in a wild population

**DOI:** 10.64898/2026.07.11.737918

**Authors:** Tomos Potter, Hanna Kokko, David N. Reznick, Joseph Travis, Beth Watson, Paul Bentzen, Ronald D. Bassar

## Abstract

If an individual’s niche is determined by its genotype, then competition for limiting resources should be most intense among individuals of the same genotype. Theory predicts this will act to maintain genetic variation, but whether this mechanism operates under natural conditions is unclear. Using long-term observations of a population of free-living Trinidadian guppies, we asked (i) whether competition was strongest between kin, and (ii) whether this process maintained genetic variation. Competition between kin was 1.5-1.8 times stronger than that between non-kin. This contributed to balancing selection: after ∼10 generations, variation was 29% higher than expected under drift. Our results show that relatedness can play a major role in structuring ecological competition, with broader consequences for theories of inclusive fitness.

**One-sentence summary:** Heritable variation is maintained due to resource competition being more intense among kin.

## Main text

Natural populations typically harbour far more genetic variation than expected under mutation-selection balance (Bonnet *et al*. 2022; Connallon & Czuppon 2026; Lewontin 1974; Ruzicka *et al*. 2026). How genetic variation is maintained in natural populations remains a fundamental question in evolutionary biology (Ayala & Campbell 1974; Charlesworth 2015; Christie & McNickle 2023; Connallon & Czuppon 2026; Gómez-Llano *et al*. 2024; Kimura & Crow 1964; Lewontin 1974; Ruzicka *et al*. 2026). One theoretical mechanism that can maintain genetic variation is balancing selection driven by negative frequency-dependence (Ayala & Campbell 1974; Christie & McNickle 2023; Ruzicka *et al*. 2026). This occurs when a genotype’s fitness is greatest when it is rare, thereby limiting its chances of fixation or extinction. Several forms of negative frequency-dependent selection have been supported by robust empirical evidence from natural populations. For example, rare genotypes have been shown to experience reduced predation or herbivory (Goldberg *et al*. 2020; Madsen *et al*. 2022; Olendorf *et al*. 2006), or to have enhanced reproductive success (Estévez *et al*. 2020; Potter *et al*. 2023; Smith 1975; Svensson *et al*. 2005). A recent meta-analysis found that the strength of negative frequency-dependent selection was strongest when driven by resource competition (Gómez-Llano *et al*. 2024), suggesting this mechanism may be key to explaining higher-than-expected levels of genetic variation observed in natural populations. However, the presence of negative frequency-dependence does not guarantee that balancing selection will occur (Ruzicka *et al*. 2026), and examples of competition-dependent balancing selection in natural systems are rare.

Although often credited to Lewontin (1974), the idea that competition for limiting resources could generate balancing selection was first described by Dobzhansky (1951) and later formalised by Levene (1953). The central idea is that an individual’s genotype determines its niche, and that each genotype occupies a subtly different niche. As such, competition for limiting resources will be most intense among individuals bearing the same genotype - an idea pithily expressed by Lewontin (1974) as “a genotype is its own worst enemy”. To ecologists, this mechanism has clear parallels with models of coexistence in which the strength of intraspecific competition must exceed that of interspecific competition for both competing species to coexist (Chesson 2000; Macarthur & Levins 1967). Despite the mechanistic similarity with established models of ecological coexistence, Lewontin doubted the likelihood of this form of balancing selection, calling it “…really a rather extreme hypothesis” (1974). Specifically, he doubted whether the necessary conditions – a sufficiently complex niche space in which each genotype is a specialist on and limited by a different resource – would be found in nature.

Indeed, empirical evidence of this type of frequency dependence typically considers competition between just two variants occupying two niches (Antonovics & Ellstrand 1984; Arnqvist *et al*. 2016; Ascensao *et al*. 2025; Kazancioǧlu & Arnqvist 2014; Kurbalija Novičić *et al*. 2020; Minter *et al*. 2015; Schluter 2003) - a far cry from the high degree of individual-level specialisation that Lewontin envisioned would be required to explain observed levels of genetic variation in natural systems. To our knowledge, evidence of this mechanism generating balancing selection under natural conditions comes from a single, wonderfully unusual study system: the scale-eating cichlid of Lake Tanganyika (Hori 1993). In this species, there are genetically-determined left- and right-mouthed morphs: their respective niches are the sides from which they attack their prey. Prey become more defensive of attacks from the side of the most abundant morph. As such, the rare morph has a fitness advantage through enhanced feeding success, and competition-mediated frequency dependence maintains the polymorphism in the population over time (Hori 1993). Whether this mechanism operates at the much finer scale of genotype-to-niche mapping envisaged by Lewontin, such that natural competitive interactions are structured by relatedness, remains unknown.

Here, we test whether competition is more intense among relatives and whether this mechanism generates balancing selection in a free-living population of Trinidadian guppies (*Poecilia reticulata*). Guppies are small, freshwater, live-bearing fish, native to the streams and rivers of Venezuela and the Caribbean island of Trinidad. In Trinidad, guppies are found in a variety of community types which can be broadly defined by the presence or absence of large predatory fish species (Deacon *et al*. 2018; Magurran 2005; Reznick *et al*. 2001; Reznick & Endler 1982). In upstream communities where predators such as the pike cichlid (*Crenicichla alta*) and wolf-fish (*Hoplias malabaricus*) are absent, guppy populations reach high densities and are strongly regulated by resource competition (Bassar *et al*. 2013; Reznick *et al*. 2001, 2019; Travis *et al*. 2014, 2023a). Analysis of stable isotopes has shown that there is considerable individual-level variation in trophic position and origin of dietary carbon in these populations (Anaya-Rojas *et al*. 2023), suggesting that individual guppies differ in their resource niche. This scenario – of intense intraspecific competition for limiting resources and individual variation in niche use – provides an ideal opportunity to ask whether (i) competition is most intense among related individuals, and (ii) whether this generates balancing selection under natural conditions.

We addressed these aims using data from a long-term mark-recapture study of guppies in a natural stream in the Northern Range mountains of Trinidad (Travis *et al*. 2014). We used a microsatellite-based pedigree to determine lineages of maternal descent (i.e. putatively distinct mitochondrial lineages) and then quantify the number of individuals of each lineage present in the population each month over a five-year period (see Methods for details). Interactions between mitochondrial haplotypes and diet influence organismal fitness in many systems (Aw *et al*. 2018; Ballard & Youngson 2015; Kazancioǧlu & Arnqvist 2014; Kurbalija Novičić *et al*. 2020; Latorre-Pellicer *et al*. 2016), implying that mitochondria, which are inherited matrilineally, may play a key role in individual dietary specialisation. In this study, we considered individuals to be kin if they share the same matriline and non-kin if they have different matrilines, and tested whether balancing selection is operating on matrilineal diversity.

To address aim (i) we quantified the competition coefficients for kin (*α*_*K*_) and non-kin (*α*_*N*_) by fitting generalized linear mixed effects models (GLMMs) for individual demographic rates of female guppies, including the number of kin and non-kin present in the population as predictors (Methods). A female guppy’s somatic growth rate and fecundity are strongly predicted by the amount of food she has consumed (Potter *et al*. 2021b; Reznick *et al*. 2004). If our estimates of *α*_*K*_ for these demographic rates are more strongly negative than those of *α*_*N*_, this would support the hypothesis that competition for resources is stronger among related individuals than between unrelated individuals. We note that if autosomal loci contribute to individual niche specialisation, our approach will underestimate the strength of competition among kin (e.g. we consider paternal half-sibs to be non-kin, although they are expected to share 25% of their autosomal alleles), making our test conservative. To address aim (ii) we performed Monte Carlo simulations that matched the monthly birth and death rates of our study population, but where individual demographic rates were independent of the matrilineal identity of individuals, i.e. were random with respect to the number of kin vs non-kin competitors present (Methods). These neutral simulations capture the potential range of changes in matrilineal diversity we would expect to see under drift given the observed changes in population size. We then compared the decline in matrilineal diversity observed in our study population with that expected under drift. Finally, we performed positive simulations — using estimates of *α*_*K*_ and *α*_*N*_ from our statistical models — to determine whether differences in the strength of competition between kin and non-kin could explain the observed maintenance of matrilineal diversity.

## Methods

### Study population and data collection

In March 2008, as part of a long-term study of evolution in the wild (Reznick & Travis 2019; Travis *et al*. 2014), we extended the range of guppies from a predator-rich region of the Guanapo river into a previously guppy-free section of an upstream tributary called Lower Lalaja, where the only other species of fish present was Hart’s killifish (*Anablepsoides hartii*), which is a competitor to and occasional predator of guppies. Thirty-eight females and thirty-eight males were introduced. Our study site forms a closed population because the ∼100m stretch of stream is bounded by barrier waterfalls at each end: there are no guppies present upstream of the site, and the lower barrier waterfall prevents colonisation by guppies from populations downstream. Since the introduction, we have performed a monthly mark-recapture study. Each month we attempted to capture all guppies larger than 14mm in standard length (measured from the tip of the snout to the posterior of the caudal peduncle) in our study population. Guppies less than this size are too small to safely capture and process. The probability of capturing an individual, given that it was alive, was 90-95% (Travis *et al*. 2023a). After capturing fish, we transported them to our lab for processing. Upon first capture, we gave each fish a unique identifying tattoo via subcutaneous coloured elastomer implants (Northwest Marine Technologies) and collected scales from the caudal peduncle for DNA extraction and the construction of the pedigree (see (López-Sepulcre *et al*. 2013 & Potter *et al*. 2021a for full details on microsatellite-based pedigree construction). At each capture, we identified and photographed each individual, and recorded its standard length, wet weight, sex, and the location in the stream where it was captured. Following processing, we returned the fish to the stream and released them at the site of their capture.

### Calculating the number of kin and non-kin competitors

As in previous lab-based studies of competition-mediated balancing selection (Arnqvist *et al*. 2016; Kazancioǧlu & Arnqvist 2014; Kurbalija Novičić *et al*. 2020), we focus on competition structured by relatedness with respect to matrilineal inheritance. Using the pedigree, we traced each individual’s maternal line to identify from which of the 38 founding females they were descended. We assumed that the founding females were unrelated to one another such that each founder represented a distinct matriline (please see the Supplementary Information for a test on the consequences of violating this assumption). We counted the number of females from each matriline alive in the population each month of the study. For each female in each month, we recorded the number of kin competitors as the total number of females with the same matriline present that month, and the number of non-kin competitors as the total number of females of different matrilines present that month. Note that we do not consider male competitors in our analyses. Male guppies have feeding rates that are an order of magnitude lower than those of females (Ventura *et al*. 2024), meaning that they have a limited impact on resource availability compared to females.

Due to genotyping failure, we could not describe maternal identity for 14% of individuals in the pedigree. These “broken links” in the pedigree meant that we could not identify the original founding matriline for those individuals, nor for any of their descendants (even if we identified the descendants’ mothers). The fraction of individuals for whom we could not identify the founding matriline increased over the 60-month duration of the study; on average we could not identify the founding matriline of 19% of individuals in a given month (Supplementary Material Figure S2A). We excluded individuals with unknown matrilines from our statistical analyses, and from counts of the number of kin/non-kin competitors present each month (please see the Supplementary Material for tests of how this influences our estimates of the kin and non-kin competition coefficients and their difference).

### Individual demographic rates

We quantified three individual-level demographic rates: monthly somatic growth rate, monthly survival, and monthly reproductive success. Because we are interested in the evolutionary consequences of competition within and between matrilines, our study focuses on demographic rates in female guppies only. We quantified monthly somatic growth rate as the change in standard length from one monthly observation to the next, divided by the number of days between measurements, then multiplied by thirty. We only quantified growth for individuals caught in consecutive months. We recorded that an individual was alive each month between its first and last capture. Because the probability of capturing a fish given that it is alive was high (90-95%), we assumed that individuals died the month after their final capture. To quantify monthly reproductive success, we used pedigree-relatedness data to identify all offspring of each female, then the month each was first observed. We assume that new recruits are around two months old at their first capture (Potter *et al*. 2023): it takes newborn guppies approximately two months to grow to 14mm standard length, i.e., the size at which we can first safely capture them. Monthly reproductive success followed an over-dispersed Poisson distribution (mean < variance), driven by zero inflation. Note that our estimate of monthly reproductive success combines the fecundity of the mother with the survival of her offspring over the first two months of their life.

Our goal was to assess how competition impacted these demographic rates. We assessed how the number of kin and non-kin present in the population at the start of the month influenced survival and somatic growth throughout the month. To assess reproductive success, we examined the impact of the number of kin and non-kin present in the month before conception (N.B. the gestation period of guppies is roughly one month). This is because guppies are lecithotrophic, meaning that the amount of resource provisioning to embryos is determined before fertilisation occurs (Reznick *et al*. 1996). As such, the maternal environment experienced prior to fertilization influences a female’s fecundity: females that receive less food can provision fewer eggs and consequently have lower fecundity (Reznick *et al*. 1996). We expect the effects of resource competition on female reproductive success to reflect the conditions experienced during the month prior to conception, i.e. the month in which she was developing and provisioning eggs that would be fertilised the following month.

### Statistical analyses

To assess whether competition among kin is more intense than among unrelated individuals, we fit linear mixed effects models of individual demographic rates as functions of the number of kin and non-kin present in the population, allowing us to estimate the kin-competition coefficient *α*_*K*_ and the non-kin competition coefficient *α*_*N*_. We fit separate models for monthly somatic growth rate, reproductive success, and survival. Because these demographic rates are size-dependent in guppies (Reznick *et al*. 2004), we included the focal individual’s size (log-transformed standard length) as a covariate in all models. We modelled random intercepts for individual ID, sampling month, and location in the stream in which an individual was observed that month. Modelling these grouping factors accounted for non-independence of data due to repeated measures on individuals over time, common environmental conditions experienced by the whole population within a month, and local conditions for individuals within a given location in the stream, respectively.

We also accounted for non-independence of observations among individuals that were likely to be competing locally with one another for resources in a given month. In a previous study on the same population, we identified “neighbourhoods” of interconnected pools and riffles within the stream, which were characterised by high movement of individuals within neighbourhoods, but low movement of individuals between neighbourhoods (see Potter et al. 2023 for details). Individuals within a neighbourhood are therefore those who are most likely to be occupying the same physical space and competing locally with one another for resources. We fit a random intercept that grouped individuals by their presence within a neighbourhood in a given month. If guppies tend to spatially cluster in groups of related individuals (a property termed “viscosity” by Hamilton (1964)) then the density of kin may simply reflect the density of local competitors, while the density of non-kin then represents the part of the population that the focal individual rarely interacts with. This would conflate relatedness with proximity, potentially biasing our estimates of kin and non-kin competition coefficients. Including this grouping factor as a random intercept corrects baseline estimates of demographic rates for this possibility.

All demographic rate models included the same predictors and random effects structures. For the model of somatic growth (right-skewed continuous data) we fit a linear model with a Student-t error structure, and allowed the variance to change as a function of log-transformed size; for reproductive success (over-dispersed count data) we fit a zero-inflated Poisson model with a log-link function, estimating a single intercept for the zero-inflation term; for the survival model (binary data), we fit a logistic regression assuming a Bernoulli distribution with a logit-link function. All models were fit in R version 4.5.2 (R Core Team 2026) in a Bayesian framework using the ‘brms’ package (Bürkner 2018). We scaled and centered the predictors for the number of kin and of non-kin by subtracting the mean population size (over the 60 months of the study) and dividing by the standard deviation of the population size. Because both kin and non-kin predictors were handled identically, the estimates for both *α*_*K*_ and *α*_*N*_ are directly proportional to per-capita effects. For each model, we used default (non-informative) priors and ran four independent MCMC chains with a burn-in of 2000 iterations, generating a total of 8000 posterior samples. Model diagnostics showed that all chains converged (all R-hat values were equal to 1.00), and that bulk and tail effective sample sizes were high (>1000), indicating that samples reliably characterized the means and variances of the posterior distribution. We validated our statistical models by performing posterior predictive checks, confirming that simulated data adequately matched the observed data distributions.

### Monte Carlo simulations

We assessed whether and how balancing selection was operating on matrilineal diversity using Monte Carlo simulations of the population dynamics. We performed two types of simulation: neutral and positive. In the neutral simulations, individual demographic rates were size-dependent but randomised with respect to matrilineal identity, thereby replicating the process of genetic drift. In the positive simulations, individual demographic rates were both size-dependent and dependent on the number of kin- and non-kin competitors present in the population, thereby replicating our proposed mechanism of balancing selection while accounting for drift. All simulations reproduced the observed population size each month. We diagnose the existence of balancing selection if the observed matrilineal diversity at the end of our study exceeded that predicted under drift. If the contrast between observed matrilineal diversity and that estimated from the neutral simulations identified balancing selection, a match between the observed matrilineal diversity and that estimated in the positive simulations demonstrate that the documented balancing selection is generated by competition structured by relatedness.

Each simulation started with 38 founding females, identified by their unique matriline and their initial size. In each month, we predicted the probability of growth, reproduction, and survival for each individual, based on the parameters from our statistical demographic rate models (neutral simulations: size-dependent only; positive simulations: size- and relatedness-structured competition-dependent, i.e. including *α*_*K*_ and *α*_*N*_). For each month of the study, we counted the total number of female recruits into the population *N*_*r*_ and the total number of female survivors *N*_*s*’_. Recall that we assume that an individual’s birth took place two months prior to its observed recruitment. Each month, we randomly selected *N*_*s*’_ individuals from the previous month’s population, with the probability of selection weighted by an individual’s size-dependent survival probability. We then selected a pool of potential breeders from the population two months earlier (i.e. females that were alive when that month’s new recruits were born) and randomly allocated the number of new recruits among these potential breeders by sampling with replacement. The survivors and new recruits together form that month’s observed population.

In our simulation, we want the distribution of reproductive success to match the zero-inflated Poisson distribution observed in the stream. Ideally, we would assign the total number of offspring produced by a single mother in a given month to a randomly selected individual in the simulation. However, because we could not identify the mother for 14% of individuals, we took a different approach: we further split the pool of potential breeders into two parts, representing the zero-inflation and Poisson parts of the distribution. We randomly assigned potential breeders into the zero-inflated group with a probability equal to the zero-inflation intercept estimate from our statistical model of reproduction. We then randomly selected *N*_*r*_ individuals (with replacement, weighted by their neutral- or positive-predicted rate) from the Poisson part of the group to identify the mothers of the new recruits. Offspring inherited their mother’s matrilineal identity and entered the population with size = 16.4 mm (the mean standard length of female guppies at first capture in our study population). To account for genotyping failure, we randomly sampled *N*_*g*_ individuals (without replacement) from the new recruits, where *N*_*g*_ was the observed number of individuals that month for whom we could not identify their mother. Offspring of these individuals inherited their mother’s unknown matriline, thereby recreating the effect of genotyping failure on our observed matrilineal diversity. At the end of the simulation we recorded matrilineal diversity as the number of unique matrilines present in the population in each month.

We ran each simulation 10,000 times, to obtain a range of values of matrilineal diversity expected under the neutral and positive scenarios. We ran a second positive simulation in which we multiplied the values of *α*_*K*_ by 2 to further test the role of kin-competition in generating balancing selection. All simulations were conducted in R version 4.5.2 (R Core Team 2026).

## Results

### Does competitor relatedness impact individual demographic rates?

Competitor relatedness affected growth and reproductive rates but not survival. The parameter estimates here are the posterior modes with the 95% credible intervals given in brackets. Monthly somatic growth rates decreased log-linearly with body size (GLMM, N = 13,880, β_*z*_ = –4.01, 95% CI [–4.11, –3.91]) and with the number of competitors present in the population that month. Specifically, the effect of competition with kin (*α*_*K*_ = –0.56 [–0.74, –0.38]) was 1.5 times that of the effect of competition with unrelated individuals (*α*_*N*_ = –0.37 [–0.49, –0.25];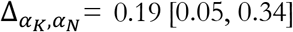; fig 1A). The monthly reproductive rate increased log-linearly with body size (GLMM, N = 17,765, β_*z*_ = 3.15 [2.86, 3.44]), and the negative effect of competition with kin (*α*_*K*_ = –1.01 [–1.43, –0.57]) was 1.8 times that of the effect of competition with non-kin (*α*_*N*_ = –0.55 [–0.71, – 0.38];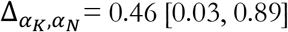; fig 1B). Monthly survival rates were not influenced by the number of related (GLMM, N = 17,765, *α*_*K*_ = 0.14 [–0.27, 0.56]) or unrelated competitors present (*α*_*N*_ = 0.07 [–0.13, 0.25]; fig1C).

**Figure 1.**
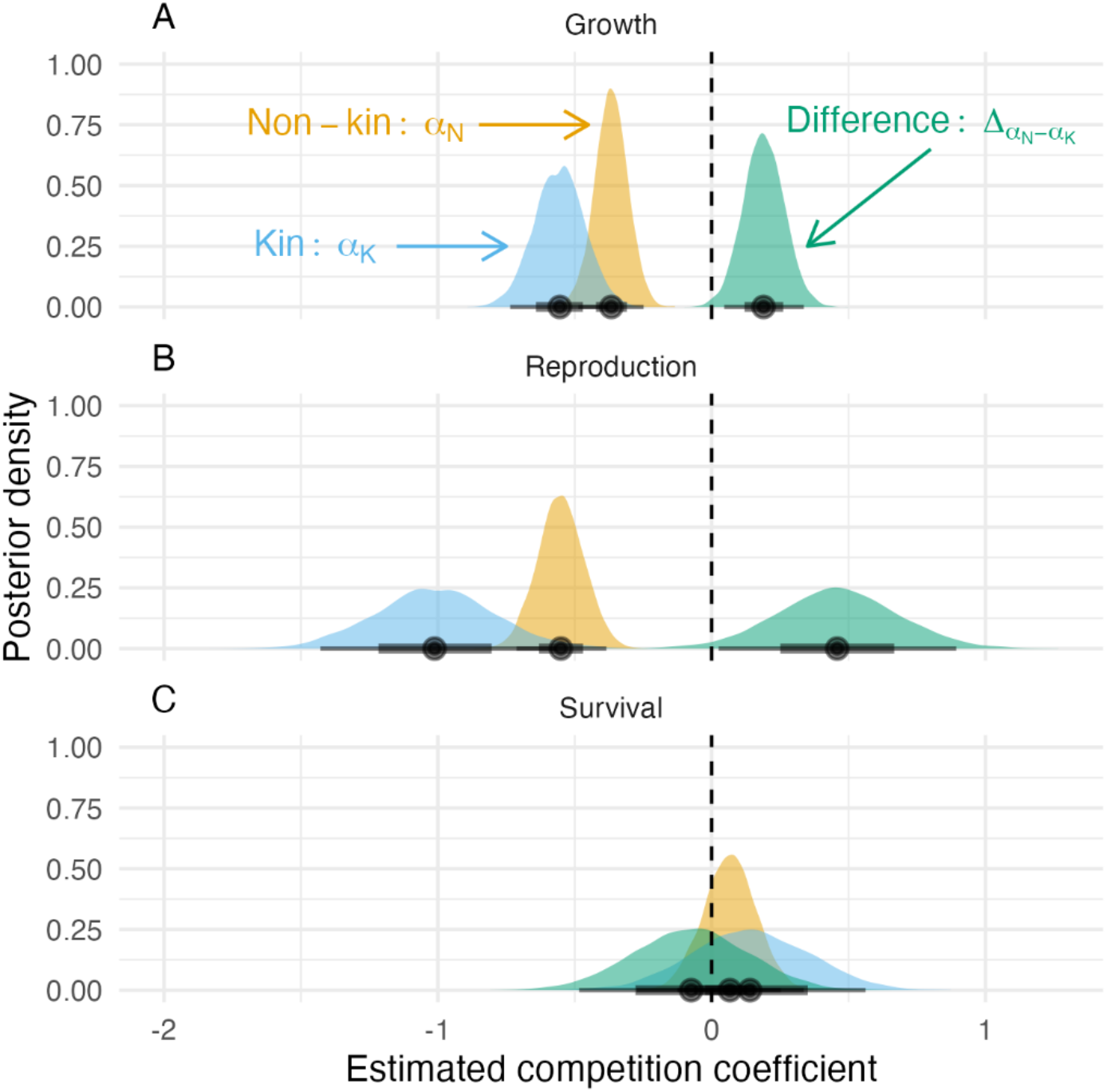
Competition among kin limits growth and reproduction, but not survival. Posterior distributions of competition coefficients for kin (*α*_*K*_, in blue) and non-kin (*α*_*N*_, yellow), and the difference between them 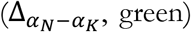, for **(A)** monthly somatic growth rate, **(B)** monthly reproductive rate, and **(C)** monthly survival probability. Black points are mean values, thick bars are 67% credible intervals, thin bars are 95% credible intervals. The x-axis gives the value of the competition coefficient (the vertical dashed line highlights zero), and the y-axis gives the density of the posterior distribution.

### Changes in diversity over time: is balancing selection operating?

While there was a steep decline in the matrilineal diversity observed at the start of the study, diversity largely stabilized thereafter (fig 2). Eighteen of the founding females died in the first month of the experiment, sixteen of which died without leaving behind any daughters. Of the thirty-eight founding females, only twenty produced daughters who were recruited into the population, thereby continuing their matrilines. The observed number of matrilines present in the population gradually decreased from 22 in the second month, to 18 by the end of the study period in month 60 (fig 2A). The modal neutral prediction matches the observed matrilineal diversity from months 20 to 38, then begins to decline relative to the observations (fig 2A). The observed number of matrilines remaining at the end of the study was 29% higher than the modal value predicted under the neutral model (modal final number of matrilines = 14): at the end of the study, the observed number of matrilines was greater than the number predicted in the null model in 97.7% of 10,000 simulations (fig 2B), suggesting that matrilineal diversity was subject to modest balancing selection.

**Figure 2.**
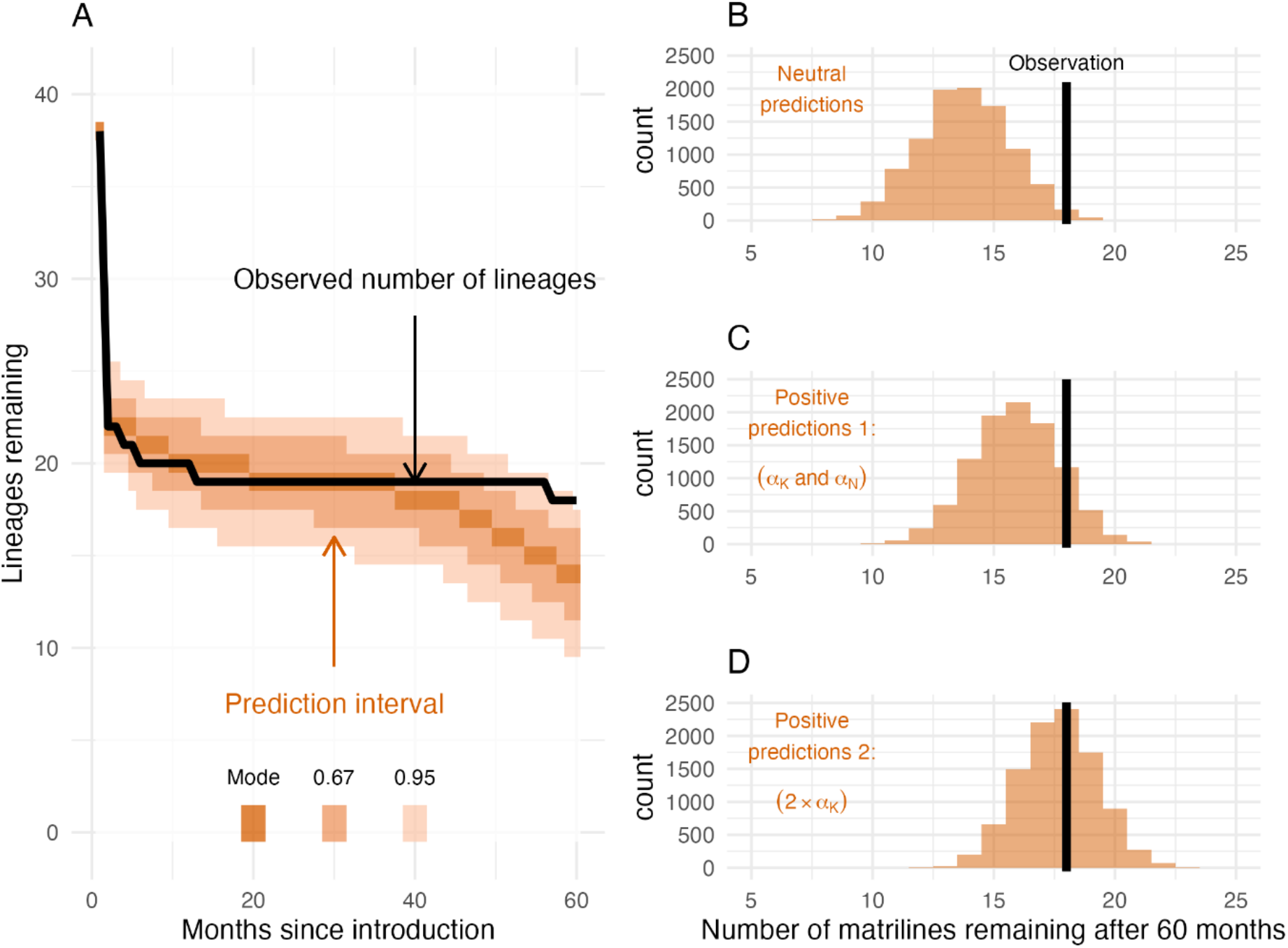
Competition among kin contributes to balancing selection. Comparing the observed change in matrilineal diversity (black lines) to those expected under three different Monte Carlo simulations. **(A)** Observed and predicted change in matrilineal diversity (y-axis) over the course of the study (x-axis) under the neutral simulation where vital rates were independent of matrilineal identity, i.e. the expectation under drift. The panels on the right show the distribution of predictions of matrilineal diversity at the end of the study (in month 60) for (B) the neutral simulations, (C) positive simulations where vital rates were dependent on the number of kin- and non-kin competitors, using the estimates of *α*_*K*_ and *α*_*K*_ obtained from our statistical models, and (D) positive simulations where *α*_*K*_ was double the value we estimated in our statistical models. All simulations tracked the observed population dynamics and were repeated 10,000 times.

In our first positive simulation we used the estimated values of *α*_*K*_ and *α*_*N*_ from our statistical models to test how competition between kin and non-kin contribute to balancing selection. In this scenario, after 60 months the mode of the predicted number of remaining matrilines was 16, which is still less than the observed number (fig 2C): the effects of kin and non-kin competition operating via growth and reproduction explained half of the difference between the null scenario and the observed data. Our second positive simulation demonstrated that competition among kin impacting reproduction and growth would need to be twice as strong for the modal prediction to match the observation (fig 2D).

## Discussion

For competition-mediated balancing selection to occur, competition must be strongest among related individuals (Dobzhansky 1951; Levene 1953; Lewontin 1974). In our study system, competition was almost twice as intense among related individuals as between unrelated individuals: to paraphrase Lewontin, guppy relatives are their own worst enemies. The two demographic rates affected by the number of kin and non-kin in the population – somatic growth and reproduction – are both highly dependent on resource intake in guppies, strongly suggesting that the underlying mechanism is limiting competition for food. Our Monte Carlo simulations demonstrate that the effects of this enhanced kin competition contributed to balancing selection, thereby maintaining matrilineal diversity at a rate higher than expected under drift. As such, our results provide evidence that competition-mediated balancing selection operates under natural conditions, although its strength was quite modest.

Our estimates of the ratio of kin (*α*_*K*_) vs non-kin (*α*_*N*_) competition coefficients were surprisingly high: 1.5 for growth, and 1.8 for reproduction. To place this in context, a meta-analysis of coexisting communities of competing plants found that the average ratio of intra- to interspecific competition coefficients was around 4 or 5 (Adler *et al*. 2018). This means that the difference in intensity of competition between kin and non-kin in our study amounts to roughly one-third to one-half of the difference between intra- and interspecific competition.

Despite this, our estimated values of *α*_*K*_ only explained half of the effects of balancing selection. What could explain this mismatch? One possibility is that competition-mediated balancing selection was operating on a component of fitness that we could not quantify, namely juvenile survival. Because neonates and juveniles are too small to safely collect from the stream, we lack data on survival over the first two months of a guppy’s life. Guppies typically produce litters ranging from three to fifteen offspring (Felmy *et al*. 2021), yet in our study the mean number of surviving offspring was ∼1.5. This suggests that juvenile mortality is high and may be a major contributor to variation in fitness. Furthermore, in guppies the effects of competition are felt most strongly by the smallest individuals (Potter *et al*. 2019). It is plausible that kin competition affecting juvenile survival could further contribute to balancing selection, although we cannot rule out other potential mechanisms operating via juvenile survival, such as frequency-dependent predation or disease risk.

In our study, we considered relatedness in the context of shared matrilineal descent. Previous laboratory studies have demonstrated balancing selection mediated by stronger competition within matrilines than between, which acted to maintain mitochondrial haplotype polymorphisms (Arnqvist *et al*. 2016; Kazancioǧlu & Arnqvist 2014; Kurbalija Novičić *et al*. 2020). A major limitation of our study is that we do not know if the founding females were all of distinct matrilines. We assumed this to be the case (as most studies of free-living pedigreed populations do), and note that, if this assumption is violated, our method underestimates the difference between kin and non-kin competition coefficients, making our estimates conservative (see Supplementary Material and fig S1).

Because we lacked mitochondrial genotypes, if we failed to identify an individual’s mother, we could not determine its matriline, nor that of any of its descendants. This missing data did slightly inflate our estimate of the difference between *α*_*K*_ and *α*_*N*_, albeit by less than 10% (Supplementary Material, fig S2). Despite these limitations, our results demonstrate that matrilineal diversity was maintained by balancing selection, and that this was generated by relatedness-structured competition in our study population.

The effects of balancing selection became apparent from month 38 of our study, which is eight months after the population density reached its peak (Travis *et al*. 2023a). This is exactly when we expect resource competition to be most intense, and so also the impact of competition-mediated negative frequency-dependence. This result aligns with earlier work demonstrating that selection on life-history traits in this population occurs only after the population has reached carrying capacity (Potter *et al*. 2021a; Reznick *et al*. 2019). Our results demonstrate that resource limitation can not only drive directional selection on life histories (Travis *et al*. 2023b), but can also generate negative frequency-dependent balancing selection. Previous work in the same population has demonstrated that male guppy colour patterns are subject to negative frequency-dependent selection, driven by female preference for mating with males baring rare patterns (Potter *et al*. 2023). Our results demonstrate that female guppies, too, are subject to negative frequency-dependent selection, driven by relatedness-structured resource competition.

Competition structured by relatedness directly links ecological and evolutionary processes, because its outcome simultaneously depends on and influences both the numerical dynamics and the distribution of heritable variation within the population. Heritable variation alters ecological interactions, which then feeds back on the success of genetic lineages. As such, this mechanism should be of particular interest in the field of eco-evolutionary dynamics (Bassar *et al*. 2021; Hairston *et al*. 2005; Pelletier *et al*. 2009), yet competition structured by relatedness has been seemingly overlooked. Quantifying this mechanism in natural populations requires data from long-term, multi-generational studies characterised by a pedigree. Many such studies exist (Bonnet *et al*. 2022), providing an opportunity to investigate how competition structured by relatedness influences eco-evolutionary dynamics in natural systems.

One field that has paid considerable attention to the consequences of competition among kin is the realm of inclusive fitness theory. However, our results contradict a key prediction - that competition should be minimised among kin (West *et al*. 2002). Instead, our results from a natural population demonstrate that competition can be elevated among kin, even after accounting for potential population viscosity (i.e. the tendency for related individuals to spatially aggregate due to limited dispersal). If kin compete more strongly than non-kin, this clearly alters predictions from inclusive fitness theory. For example, Taylor’s (1992) classic model demonstrated that the altruism-favouring effects of viscosity are perfectly cancelled out by the altruism-opposing effects of kin competition in a meta-population. This influential result – which decouples dispersal from kin competition - holds because of the implicit assumption of equally strong competition between kin and non-kin; our results demonstrate that this assumption warrants closer investigation.

## Acknowledgments

We thank the many field managers and interns, past and present, for their contributions to the Guppy Project, and Mahase Ramlal and family for hosting us in Trinidad.

## Author contributions

Conceptualization: TP, RDB, DNR, JT; Methodology: all authors; Investigation: TP, RDB, JT, DNR; Visualization: TP; Funding acquisition: DNR, JT, RDB, HK; Project administration: DNR, JT, RDB; Supervision: DNR, JT, HK; Writing – original draft: TP; Writing – review & editing: all authors.

## Data and materials availability

All data and code required to reproduce the analyses presented here will be posted in an online repository on acceptance.

## Ethics

All animals were collected and handled with the approval of the University of California Riverside IACUC AUP (no. a-20080008).

## Supplementary material

### If founders are related, how will this bias our estimates of competition coefficients?

Prior work has shown that there are strong density-dependent effects on growth and reproduction in guppies (Bassar *et al*. 2013; Travis *et al*. 2023a), that is, the intraspecific competition coefficient (including the combined effects of both kin and non-kin) is negative. Our goal is to determine whether kin compete more strongly than non-kin, i.e. is the kin competition coefficient *α*_*K*_ more strongly negative than the non-kin competition coefficient *α*_*N*_? To do so, we decompose monthly population density into the total number of kin and non-kin present, with respect to each individual, then use these values as predictors in models of individual demographic rates (see Methods in main text for details).

As is typical in quantitative genetic studies of wild populations, we assumed that the founding members of the population were unrelated. Specifically, we assumed that each of the founding females represented a different and distinct matriline. Because matrilineal identity is the basis of our measure of relatedness, it is important that we consider the potential consequences of this assumption for our estimates of *α*_*K*_ and *α*_*N*_. We know that there cannot be more matrilines present in the population than the number of founding females, but there may be fewer if some of the founders were e.g. sisters or matrilineal cousins. If so, this would mean that we would have systematically undercounted the number of kin and overcounted the number of non-kin with respect to individuals descended from a founding female from a non-unique matriline (i.e. who is related to other founders).

We tested how this might bias our estimates of *α*_*K*_ and *α*_*N*_ by performing a simulation study where the assumed number of lineages was greater than the true number of lineages. Each simulation began with 20 assumed lineages. The number of individuals of each assumed lineage was randomly drawn from a uniform distribution (min = 5, max = 25). We then randomly assigned assumed lineages into a smaller number of groups, reflecting the “true” lineages (groups ranged from 2 to 19 “true” lineages). This simulates scenarios ranging from a single pair of related founders (i.e. 19 lineages) to all founders belonging to one of two lineages. For each scenario, we calculated the “true” number of kin 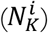 and non-kin 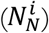 present, then calculated values of a trait *y* given the true number of lineages *i* as 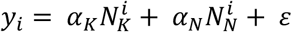, where ε is an error term randomly drawn from a normal distribution with a mean of 0 and a standard distribution of 1. We tested scenarios where competition was stronger among kin (*α*_*K*_ = −1; *α*_*N*_ = −0.5), and where competition was weaker among kin (*α*_*K*_ = −0.5; *α*_*N*_ = −1).

We then fit linear models to predict *y*_1_ as a function of the assumed number of kin 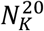 and non-kin 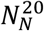. We assessed bias as the difference between the values of *α*_*K*_ and *α*_*N*_ used to simulate the data, and the estimated values of *α*_*K*_ and *α*_*N*_ returned from the linear models. Note that for both *α*_*K*_ and *α*_*N*_ to be identifiable, there must be variation in total population size between some samples. As such, we generated 5 random populations for each scenario, which differed in total population size. For each scenario of the relative strength of competition (*α*_*K*_> *α*_*N*_; *α*_*K*_< *α*_*N*_), we ran 1000 simulations and we report the mean values of estimates of *α*_*K*_ and *α*_*N*_.

We found that the assumption of unrelated founders did not bias estimates of *α*_*K*_, but systematically biased estimates of *α*_*N*_ (fig S1A). Specifically, as the number of true lineages decreased, estimates of *α*_*N*_ were increasingly biased towards the true value of *α*_*K*_, such that the true effect size (i.e. the absolute difference between *α*_*K*_ and *α*_*N*_) was increasingly underestimated (fig S1B). As such if our assumption that founders are unrelated is violated, we would have systematically underestimated the true difference between *α*_*K*_ and *α*_*N*_, i.e. our results would be conservative.

### How does genotyping failure bias our estimates of competition coefficients?

In our focal pedigree, genotyping failure meant that we could not identify the mother of 14% of individuals, which consequently meant that, on average, we could not identify the matriline of 19% of individuals in the population in a given month (fig S2A). The question then is how to handle these missing data in our analyses to minimise bias in our estimates of *α*_*K*_ and *α*_*N*_ and, crucially, their difference.

We consider two potential options: the first is to exclude individuals with unidentified matrilines from counts of competitors; the second is to count them as non-kin competitors. The first option ignores the presence of these individuals in the population, while the second treats them as though they are unrelated to any other individual. We tested the consequences of these choices by simulating data across a range of values of missing matriline identity (0%-50%) then fitting models to the simulated data to estimate *α*_*K*_ and *α*_*N*_ following the two strategies described above. In each simulation, we began with 20 lineages, then assigned the number of individuals to each lineage by drawing from a uniform distribution (min = 5, max = 25). We predicted the “true” values of a trait *y* for each individual in the same manner as in the previous section (i.e. 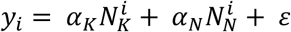). We then randomly selected a percentage of individuals from the population to represent those with unknown matrilines. As in the previous simulation, we generated data for five populations differing in total population size for each simulation run. We then fit linear models to estimate the values of *α*_*K*_ and *α*_*N*_ under the two strategies for handling unknown matrilines. Again, we considered two scenarios differing in the relative strength of kin competition (*α*_*K*_> *α*_*N*_; *α*_*K*_< *α*_*N*_), and we ran 1000 simulations and report the mean values of estimates of *α*_*K*_ and *α*_*N*_.

For scenarios where competition is stronger among kin (*α*_*K*_< *α*_*N*_), option 1 (excluding individuals with unknown matrilines from the analysis) resulted in overestimating the strength of competition both among kin and between non-kin, i.e. *α*_*K*_ and *α*_*N*_ were both increasingly biased downwards away from the true values as the proportion of individuals with unknown matriline increased (fig S2B). Using option 2 (treating unknown matrilines as unrelated competitors), estimates of *α*_*N*_ were unbiased but *α*_*K*_ was increasingly overestimated as the proportion of unknown matrilines increased (fig S2B). If competition is stronger among unrelated individuals than among kin, option 1 again results in increasing overestimation of *α*_*K*_ and *α*_*N*_, while option 2 again results in unbiased estimates of *α*_*N*_, but increasingly underestimates *α*_*K*_ relative to the true value (fig S2C). Effect sizes (i.e. the difference between *α*_*K*_ and *α*_*N*_) were increasingly overestimated with increasing percentage of unknown matrilines (figs S2D & E). However, for our observed mean value of unknown matrilines (19%), this bias is relatively small (∼10%), with option 1 minimising bias when competition is strongest among kin (fig S2D), and option 2 minimising bias when competition is strongest among non-kin (fig S2E).

## Supplementary Figures

**Figure S1.**
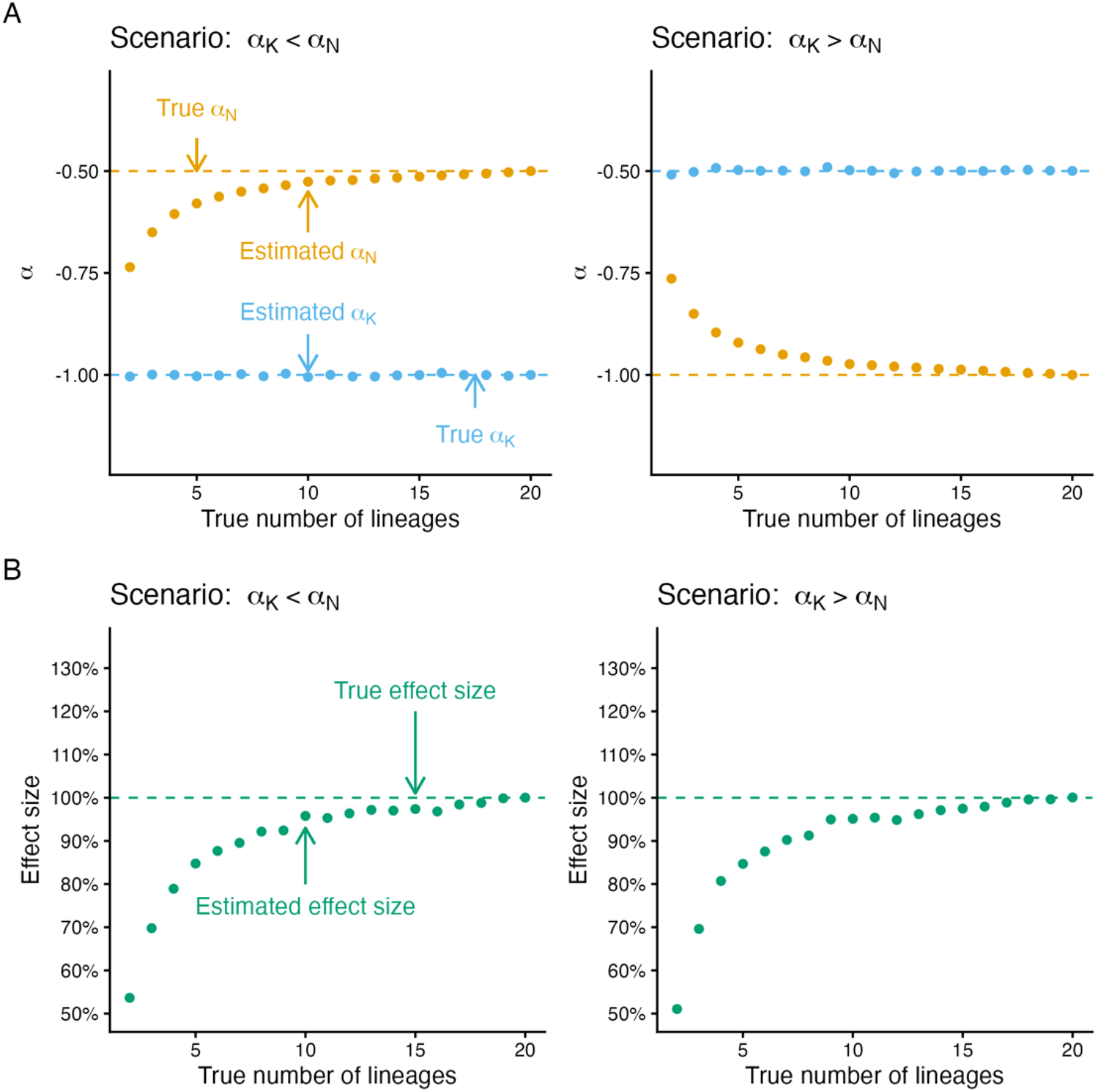
How are estimates of competition coefficients biased if the assumption that founding individuals are unrelated is violated? We simulated data based on the true number of lineages (x-axis in all plots), then estimated the kin competition coefficient *α*_*K*_ and the non-kin competition coefficient *α*_*N*_ assuming that all founding individuals were unrelated. In [A], points are estimated values of *α*_*K*_ (blue) and *α*_*N*_ (yellow), and dashed lines show the actual values used to generate the data in two different scenarios (*α*_*K*_< *α*_*N*_ and *α*_*K*_>*α*_*N*_). In [B], points are estimated effect sizes (i.e. absolute difference between *α*_*K*_ and *α*_*N*_), and the dashed line shows the actual effect size used to generate the data, for the same two scenarios. In all panels, points are mean values based on 1000 simulations of each scenario.

**Figure S2.**
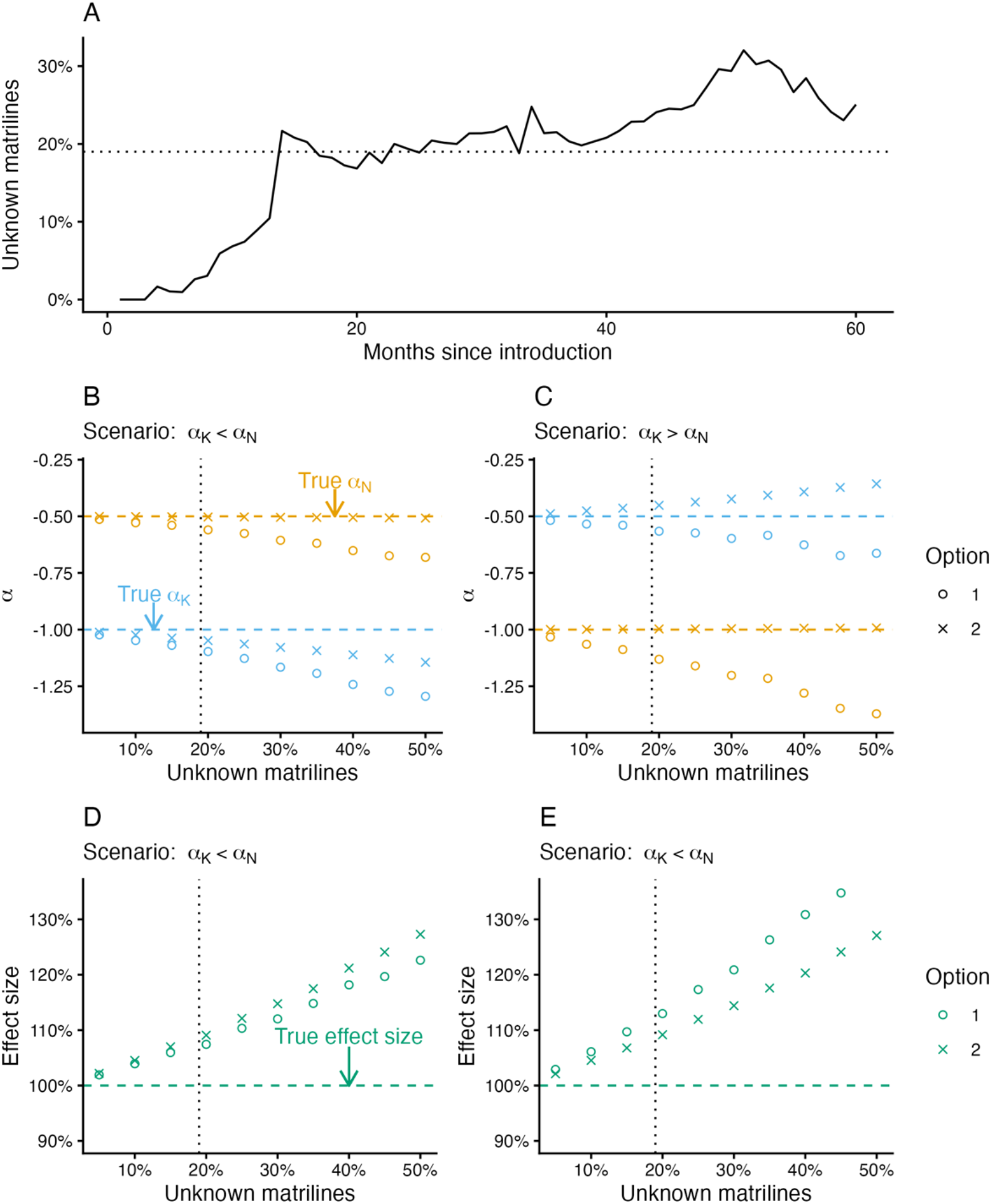
Handling data when matrilines are unknown for some individuals. (A) The observed percentage of individuals within the population with unknown matrilines (y-axis) increased over the duration of the study (x-axis). The horizontal dotted bar highlights the mean percentage of individuals of unknown matriline. We simulated data to test how model estimates of *α*_*K*_ (blue) and *α*_*N*_ (yellow) were biased when a random fraction of individuals had unknown matrilines (x-axis in panels B-E), under data handling option 1 (circles) and option 2 (crosses) for two scenarios: (B) *α*_*K*_> *α*_*N*_, and (C) *α*_*K*_> *α*_*N*_. Dashed horizontal lines show the true values of *α*_*K*_ and *α*_*N*_ used to simulate the data. Predicted effect sizes are shown for (B) *α*_*K*_> *α*_*N*_, and (C) *α*_*K*_> *α*_*N*_; true effect sizes are indicated by the horizontal dashed line. In panels B-E, the vertical dotted line highlights the observed mean percentage of unknown matrilines in our actual dataset. Points are mean values based on 1000 simulations of each scenario.

